# Melanocytes determine angiogenesis gene expression across human tissues

**DOI:** 10.1101/2020.10.26.355792

**Authors:** Shirly Freilikhman, Marianna Halasi, Alal Eran, Irit Adini

**Affiliations:** Department of Life Sciences, Ben Gurion University, Beersheva 84105, Israel; Department of Surgery, Harvard Medical School, The Center for Engineering in Medicine, Massachusetts General Hospital, Boston, MA 02114, USA; Computational Health Informatics Program, Boston Children’s Hospital, Boston, MA 02115, USA

**Keywords:** Angiogenesis pathways, APOE, FGFR1, MC1R, pigmentation, VEGFA

## Abstract

Several angiogenesis-dependent diseases, including age-related macular degeneration and infantile hemangioma, display differential prevalence among Black, as compared to White individuals. Although socioeconomic status and genetic architecture have been suggested as explaining these differences, we have recently shown that pigment production *per se* might be involved. For example, we have shown that the extracellular protein fibromodulin is a pro-angiogenic factor highly secreted by melanocytes in White but not Black individuals. Still, additional pigment-dependent angiogenic factors and their molecular mechanisms remain to be identified. Understanding the contribution of pigmentation to angiogenesis in health and disease is essential for precision medicine of angiogenesis-dependent diseases with racial disparity. Toward that goal, we compared the transcriptomes of Black and White individuals in three tissues with angiogenic activity, namely artery, whole blood, and skin. We identified several differentially expressed angiogenesis pathways, including artery morphogenesis, regulation of endothelial cell chemotaxis, and cellular response to vascular endothelial growth factor stimulus. We then demonstrated that the expression of key genes in these pathways is directly modulated by the degree of pigmentation. We further identified the precise pigment production pathway controlling the expression of these genes, namely melanocortin 1 receptor (MC1R) signaling. These results demonstrate pigment-mediated regulation of angiogenesis-related pathways and their driver genes across human tissues.

## Introduction

An enormous amount of epidemiological evidence supports a correlation between the prevalence of angiogenesis-related disease and skin color. For example, age-related macular degeneration is 55% less common in Black Americans and 46% less common in Asian Americans than in White Americans [1]. Likewise, White individuals are 1.5-fold more likely to present infantile hemangiomas than are Blacks [2]. Similarly, rosacea is 3.3-3.8 more common in White, as compared to Black individuals [3]. Yet, the molecular bases for the disparities in these and other angiogenesis-related disease prevalence between White and Black individuals remain largely unknown.

In skin, pigment-producing melanocytes are located in the basal layer of the epidermis, specifically within the extracellular matrix (ECM) that provides structural and biochemical support to surrounding cells. Melanocytes are also found in the eye, the inner ear, the heart, and most connective tissues, such as bone and meninges [4–6]. While Black and White individuals possess similar numbers of melanocytes, differences in melanocyte structure and function, as well as in the degree of pigment production, have been reported [7]. In addition to producing pigment, melanocytes can also affect gene expression and related functions of other tissues via secreted proteins [8–11]. Recently, we described a new paradigm wherein melanocytes regulate angiogenesis via secretion of fibromodulin (FMOD), an ECM protein belonging to the small connective tissue proteoglycan family [12]. We found that low pigment-producing melanocytes express high levels of FMOD, which, in turn, tilts the angiogenic balance towards the initiation of angiogenesis. In contrast, highly pigmented melanocytes express low levels of FMOD, leading to reduced angiogenic capacity. Secreted FMOD regulates angiogenesis in the microenvironment through its impact on TGF-β1 secretion [11]. However, these and other angiogenesis-related processes modulated by pigment levels remain to be discovered. Moreover, it is still not known which differences in angiogenesis-related gene expression seen between Black and White individuals can be attributed to differential pigment levels, as opposed, for example, to common genomic variation. Understanding pigment-dependent gene expression is, therefore, essential for elucidating the molecular mechanisms underlying angiogenesis-related diseases characterized by racial disparity, and ultimately adopting precision care approaches for their management.

In the present study, we compared the transcriptomes of Black and White individuals in three tissues with angiogenic activity, namely artery, whole blood, and skin. We identified several differentially expressed angiogenesis pathways, including artery morphogenesis, regulation of endothelial cell chemotaxis, and cellular response to vascular endothelial growth factor stimulus. We further demonstrated that differentially expressed pathway genes, including *VEGFA*, *APOE*, and *FGFR,* are expressed by mouse melanocytes and mouse melanoma cells, where their levels are directly modulated by the degree of pigmentation. Finally, we identified the precise pigment production pathway controlling the expression of these genes, namely melanocortin 1 receptor (MC1R) signaling. These results demonstrate pigment-mediated regulation of angiogenesis-related pathways and their driver genes across human tissues. Based on these findings, we propose that modulation of these genes or their products may be leveraged for precision therapeutics of angiogenesis-dependent diseases characterized by racial disparity.

## Materials and methods

### Human RNAseq data analysis

To identify angiogenesis-related genes and pathways that might be regulated by pigmentation in humans, we analyzed RNAseq data from the Genotype-Tissue Expression (GTEx) project, under authorized usage (dbGaP accession phs000424.v7.p2) [13]. Specifically, we focused on samples from Black and White individuals and tissues with angiogenic activity shown to be regulated by melanocyte secretions [10,11]. We compared gene expression between Black and White individuals in non-sun-exposed skin (257 samples from 35 Black and 222 White individuals), artery (336 samples from 50 Black and 286 White individuals), and whole blood (815 samples from 112 Black and 703 White individuals).

FastQC [14] version 0.11.6 was used to ensure the quality of the RNAseq data, followed by quality-trimming using Trimmomatic [15] version 0.36, and alignment to the GRCh38.p12 reference human genome using STAR version 2.5.3b [16], focusing on uniquely mapping reads. Genecode V29 genes were then quantified using HTSeq-count [17] version 0.11.1 and filtered to include only those with more than one count per million (CPM) in at least 100 samples. Finally, trimmed mean of M-values (TMM) normalization [18] was applied using edgeR version 3.22.5 in R version 3.5.2.

### Differential gene expression analysis

Limma [19] version 3.36.5 was used to compare gene expression between samples from Black and White individuals. Age, gender, postmortem interval, and RNA integrity were used as covariates in tissue-specific linear models.

### Gene set enrichment analysis (GSEA)

GSEA [20] was used to assess pathway- and mechanism-level differential expression between samples from Black and White individuals.

### Network level analyses

The STRING database of protein-protein interactions (PPIs) was used for all network level analyses and visualizations [21].

### Graphics

All plots were generated in ggplot2 [22] version 3.3.0.

### Cell culture

Mouse melanocytes, specifically Melan-a cells from black C57BL/6J mice and Melan-e1 cells from white C57BL/6J Mc1r−/− mice [23] (gift from Prof. Dorothy C. Bennett, University of London), were grown in RPMI-1640 medium (Sigma) supplemented with 10% fetal bovine serum (Peak Serum), 1% penicillin-streptomycin (GIBCO), 0.5% 2-mercaptoethanol (GIBCO) and 1% HEPES (GIBCO) at 37°C in a 10% CO_2_-containing environment. The Melan-a cells were supplemented with 200 nnM tetradecanoyl phorbol acetate (TPA) (Sigma), while the Melan-e1 cells were supplemented with 200 nM TPA and 40 pM cholera toxin (Sigma). B16-F1 melanoma cells from black C57BL/6J mice [24] and B16-4GF melanoma cells from white C57BL/6J Mc1r−/− mice (gift from Prof. Hyejung Jung, Ewha Womans University, Seoul, Korea) were grown in DMEM (GIBCO) supplemented with 10% fetal bovine serum and 1% penicillin-streptomycin at 37°C, in a 5% CO_2_-containing environment.

### Total RNA extraction and quantitative real-time PCR (qRT-PCR)

Total RNA was extracted in triplicate using the IBI Isolate Total Extraction Reagent System (IBI Scientific), and cDNA was synthesized using a High Capacity cDNA Reverse Transcription Kit (Applied Biosystems). qRT-PCR was performed to determine relative gene expression in pigmented versus non-pigmented mouse melanocytes using a LightCycler 480 instrument (Roche) with PrimeTime Gene Expression Master Mix (IDT DNA) and the following primers: Apoe–S, TGGAGGCTAAGGACTTGTTTC; Apoe-AS, CACTCGAGCTGATCTGTCAC; Fgfr1–S, GAGCATCAACCACACCTACC; Fgfr1-AS, CCTTACACATGAACTCCACATTG; Vegfa-S, AGAAAGACAGAACAAAGCCAGA; Vegfa-AS, TGGTGACATGGTTAATCGGT; Ppia–S, CAAACACAAACGGTTCCCAG; and Ppia-AS, TTCACCTTCCCAAAGACCAC. Expression levels were measured relative to that of *Ppia*, encoding peptidylprolyl isomerase A. Statistical significance was assessed using t-tests.

### Overall statistical considerations

All P values were corrected for multiple testing using the Benjamini-Hochberg approach [25], ensuring that the false discovery rate (FDR) of this study is below 0.05.

## Results

### Angiogenesis pathways are differentially expressed between Black and White individuals

We first sought to identify genes and pathways that are robustly differentially expressed between Black and White individuals in tissues with angiogenic activity shown to be regulated by melanocyte secretions [10,11]. Toward this goal, we leveraged large-scale RNAseq data of the GTEx project, specifically that from blood, artery, and skin that had not been sun-exposed. We detected 700 differentially expressed genes (DEGs) in whole blood, 1051 DEGs in the tibial artery, and 1301 DEGs in non-sun-exposed skin, at FDR ≤ 0.05 (**ESM 1**). Gene set enrichment analysis was next used to identify gene expression differences between White and Black individuals at the whole pathway level. Several angiogenesis pathways were found to be up-regulated in White individuals, including artery morphogenesis (P = 2.10 × 10^−3^), regulation of endothelial cell chemotaxis (P = 3.99 × 10^−3^), and cellular response to vascular endothelial growth factor stimulus (P = 7.91 × 10^−3^) (**Fig. 1a**). These findings are consistent with the observed elevated burden of angiogenesis-dependent diseases in White, as compared to Black individuals [1–3].

**Fig. 1.**
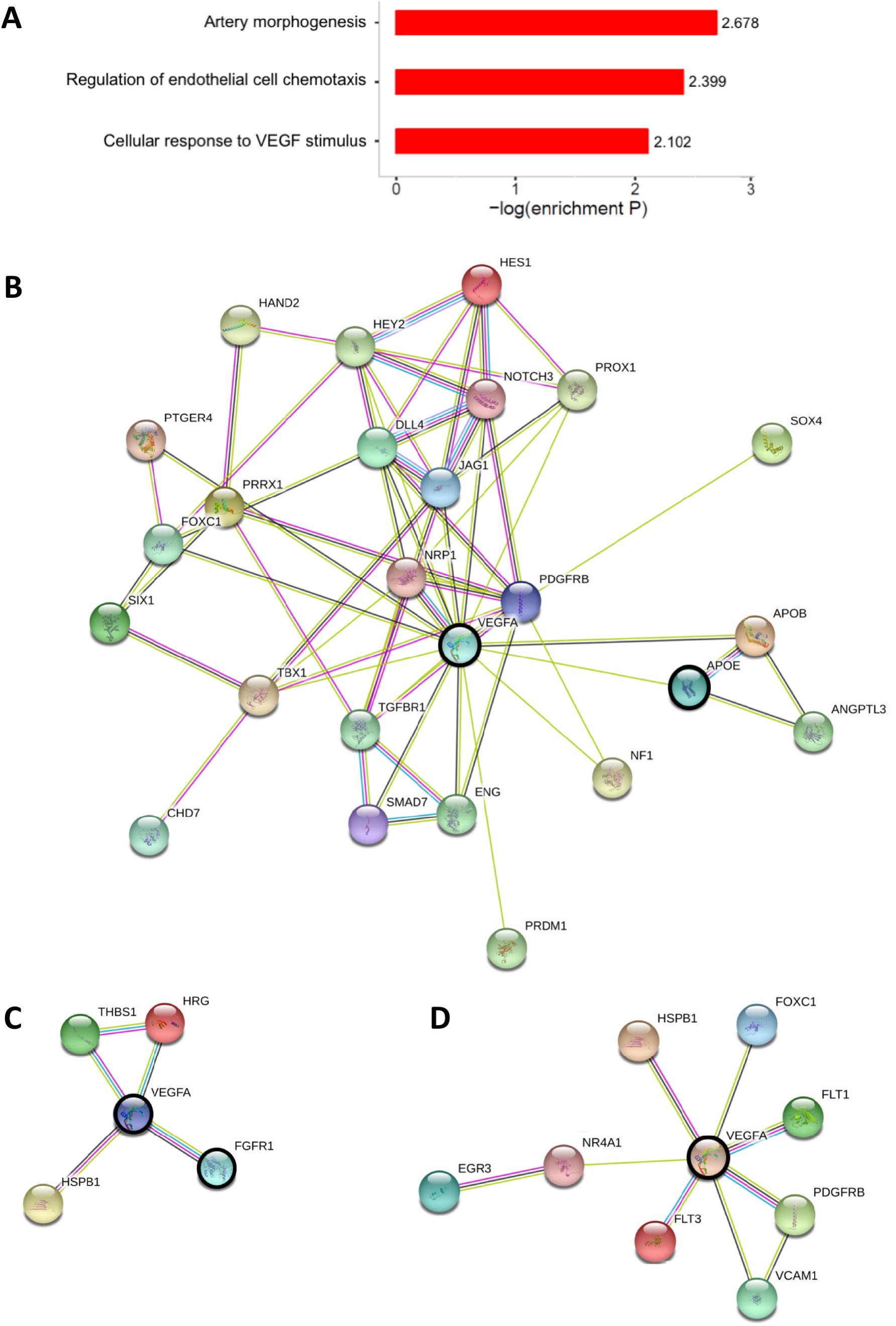
Angiogenesis pathways differentially expressed between Black and White individuals and the network structure of interactions between their leading-edge gene products. (**a**) Convergence of DEGs between Black and White individuals on angiogenic functions. Shown are enriched Gene Ontology (GO) annotations among DEGs between 112 black and 703 white individuals, as measured in whole blood RNAseq. (**b-d**) Relationships between the protein products of genes driving the up-regulation of (**b**) artery morphogenesis, (**c**) regulation of endothelial cell chemotaxis, and (**d**) cellular response to VEGF stimulus in Black, as compared to White individuals. Shown is the PPI network structure of products of leading-edge genes, namely those driving pathway-level differential expression between Black and White individuals. Nodes represent proteins, while their interactions are denoted by color-coded edges: Magenta, experimentally determined interactions; cyan, manually curated interactions; green, interactions reported in PubMed abstracts; black, co-expression; and blue, protein homology. VEGFA, APOE, and FGFR1 are highlighted by a bold black circle because of their network centrality and implication in diseases of racial disparity.

Subsequent network analyses identified several hub genes driving the differential expression of these pathways (**Fig. 1b-d**). For example, leading-edge genes (i.e. whose expression correlates most with skin color) of the artery morphogenesis pathway, whose protein products are most central to the PPI network of this pathway, include those encoding vascular endothelial growth factor A (VEGFA), platelet-derived growth factor receptor beta (PDGFRB), Jagged-1 (JAG1), Notch receptor 3 (NOTCH3), and Hes family BHLH transcription factor 1 (HES1). Notably, apolipoprotein E (APOE), which is directly connected to the products of three other members of this network, is known for its anti-angiogenic functions [26] and has been implicated in several diseases with racial disparity, including Alzheimer’s disease [27] and age-related hearing loss [28] (**Fig. 1b**). Similarly, **Fig. 1c** depicts the PPI network of leading-edge gene products involved in endothelial cell chemotaxis, highlighting the centrality of VEGFA. One notable member of this network is fibroblast growth factor receptor 1 (FGFR1), which was shown to be a key driver of melanoma angiogenesis and treatment resistance [29], common phenomena in Black individuals [30]. Finally, interactions between the protein products of leading-edge genes of the cellular response to the VEGF stimulus pathway are shown in **Fig. 1d.**

### *VEGFA, APOE*, and *FGFR1* are differentially expressed between Black and White individuals

We next examined the expression of genes driving the observed pathway-level differential expression between Black and White individuals, specifically those genes whose protein products are central to the PPI network of these pathways (designated by bold circles in Fig. 1b-d). We found that *VEGFA* expression is down-regulated in Black, as compared to White individuals (log fold-change (FC) = 1.49, adjusted P = 3.96 × 10^−4^; **Fig. 2a**), while *APOE* and *FGFR1* are up-regulated in Black, as compared to White individuals (*APOE* FC = 2.29, adjusted P = 1.72 × 10^−7^; **Fig. 2b**; *FGFR1* FC = 1.36, adjusted P = 8.60 × 10^−3^; **Fig. 2c**). To test whether the expression of these genes directly depends on pigmentation levels, we next considered two cellular systems of melanocortin 1 receptor (MC1R) signaling, a major determinant of human pigmentation.

**Fig. 2.**
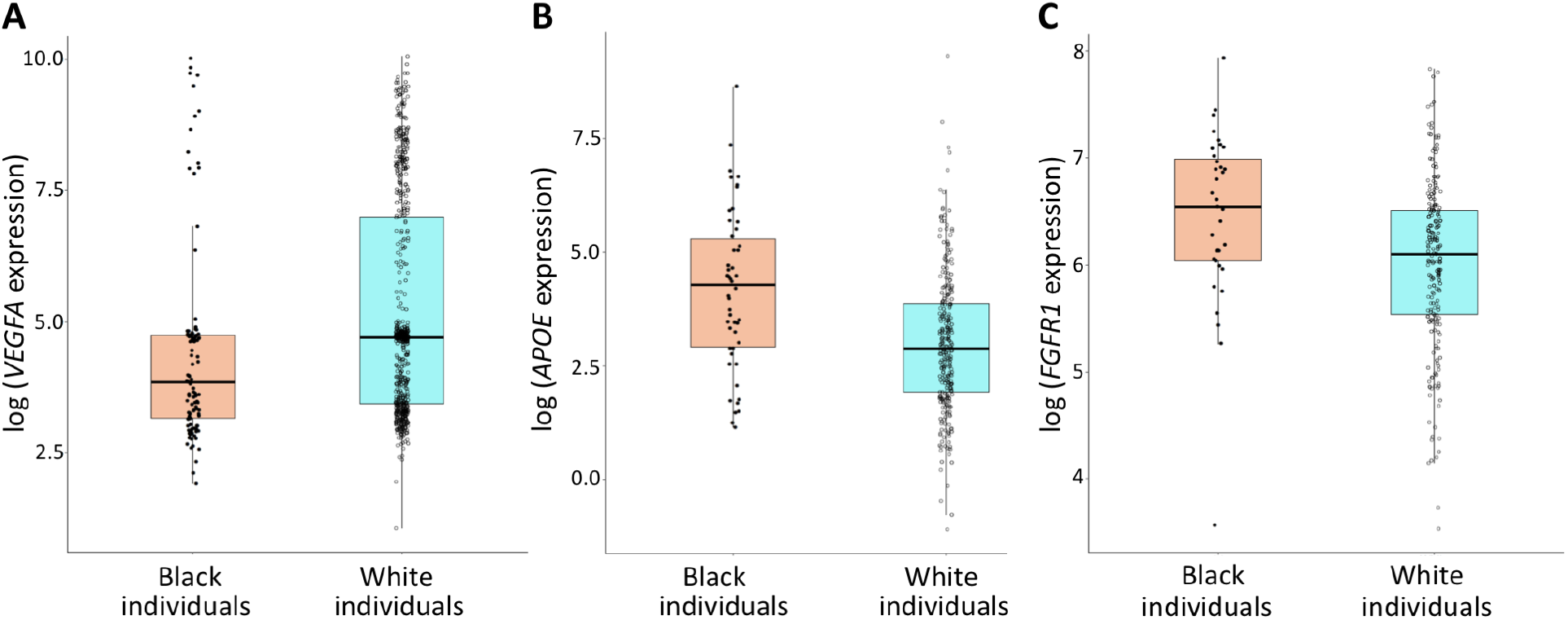
Differential expression of *VEGFA*, *APOE*, and *FGFR1* between Black and White individuals. (**A**) Down-regulation of *VEGFA* expression in whole blood from 112 Black, as compared to 703 White individuals. (**B**) Up-regulation of arterial *APOE* expression in 50 Black, as compared to 286 White individuals. (**C**) Up-regulation of *FGFR1* expression in non-sun-exposed skin from 35 Black, as compared to 222 White individuals.

### *Vegfa, Apoe*, and *Fgfr1* expression is pigmentation-dependent and controlled by melanocortin 1 receptor (Mc1r) signaling

MC1R is a key regulator of pigment production, whose genomic variants are largely responsible for pigmentation differences in humans [31]. Therefore, we used two cellular systems of murine MC1R function to examine the impact of pigmentation on *Vegfa*, *Apoe*, and *Fgfr1* expression. The first is an isogenic system of *MC1R* loss-of-function in mouse melanocytes [23], while the second is an isogenic system of *MC1R* loss-of-function in the mouse B16 melanoma cell line [32]. Each system allows for analysis of the direct effect of MC1R function, and thereby pigment production, on gene expression in an isogenic background. Using these two systems, we demonstrated that the expression of *Vegfa*, *Apoe*, and *Fgfr1*, which drive the differential expression of angiogenesis pathways between White and Black individuals, depends on pigment production via MC1R. Specifically, we found that *Vegfa* gene expression is 1.56-fold up-regulated in non-pigmented, as compared to pigmented mouse melanocytes (P = 6.58 × 10^−3^), and 5.12-fold up-regulated in non-pigmented, relative to pigmented, mouse melanoma cells (P = 1.43 × 10^−6^; **Fig. 3a**). Moreover, *Apoe* levels were 235.56-fold up-regulated in pigmented mouse melanocytes, as compared to their non-pigmented counterparts (P = 8.61 × 10^−3^), and 5.05-fold up-regulated in pigmented versus non-pigmented mouse melanoma cells (P = 1.79 × 10^−2^; **Fig. 3b**). Similarly, *Fgfr1* levels were 4.5-fold up-regulated in pigmented versus non-pigmented mouse melanocytes (P = 1.32 × 10^−2^), and 111.6-fold up-regulated in pigmented mouse melanoma cells, relative to non-pigmented such cells (P = 1.47 × 10^−2^; **Fig. 3c**). These results suggest that the differential expression of angiogenesis pathways between Black and White individuals depends on pigmentation levels and is regulated by MC1R signaling.

**Fig. 3.**
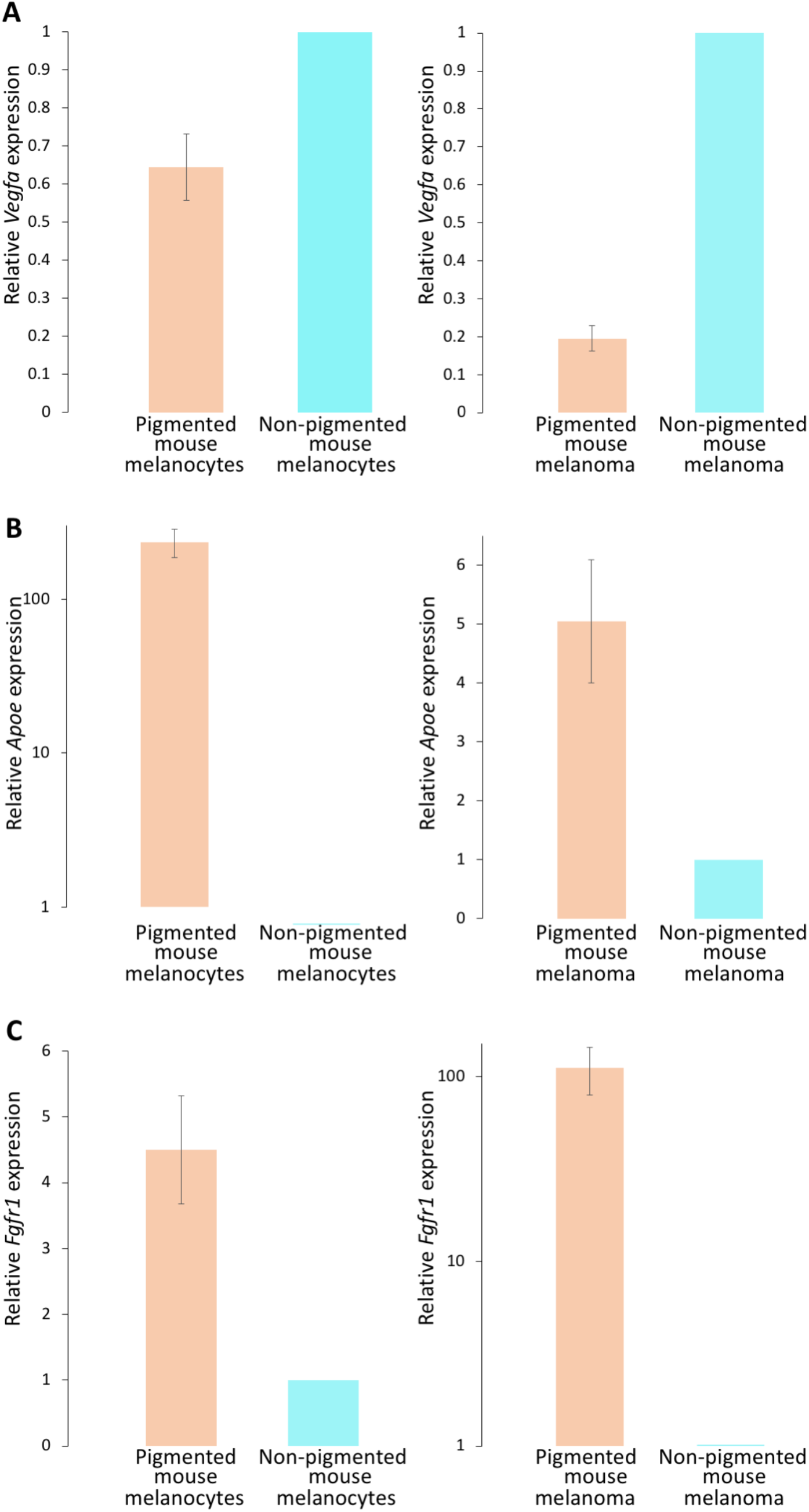
MC1R-dependent expression of *Vegfa*, *Apoe*, and *Fgfr1.* Two isogenic cellular systems of MC1R loss-of-function were used to define relationships between pigmentation and the expression of (**a**) *Vegfa*, (**b**) *Apoe*, and (**c**) *Fgfr1*. Consistent with findings in Black and White individuals shown in Fig. 2, *Vegfa* was found to be down-regulated in pigmented, as compared to non-pigmented cells, while *Apoe* and *Fgfr1* were both up-regulated in pigmented versus non-pigmented cells.

## Discussion

This study identified angiogenesis pathways that are differentially expressed between Black and White individuals, as well as their driver genes. It was subsequently shown that the expression of these driver genes, namely *VEGFA*, *APOE*, and *FGFR1*, depends on MC1R signaling and is thereby directly modulated by pigmentation levels. As such, this study improves our understanding of the impact of pigmentation on angiogenesis, and moves us one step closer toward precision medicine of angiogenesis-dependent diseases characterized by racial disparity.

Melanogenesis, the complex process by which melanocytes produce melanin, is stimulated by several factors, including ultraviolet (UV) irradiation, melanocyte-stimulating hormone (MSH), fibroblast growth factor 2 (FGF2), and the tyrosinase-catalyzed oxidation of tyrosine. The resulting pigmentation is generally classified either by presence of the black-brown eumelanin pigment that acts as a photoprotective anti-oxidant, or the yellow to reddish pheomelanin that serves as a phototoxic pro-oxidant [33]. Upon UV irradiation, α-MSH secreted by keratinocytes binds to MC1R, a G protein-coupled receptor expressed on the melanocyte surface that serves as a key regulator of pigment production, controlling whether eumelanin or pheomelanin is produced in the hair and skin. In mouse models and human studies, polymorphisms in *MC1R* have been associated with the levels of eumelanin and pheomelanin [34–36]. The present study suggests that MC1R function controls the expression of angiogenesis pathways by directly modulating the expression of their driver genes. Consistent with the present results showing up-regulation of angiogenesis pathways in White, as compared to Black individuals, we have previously shown that non-pigmented melanocytes promote angiogenesis *in vitro* and *in vivo,* and that FMOD contributes to this effect [11]. The present study thus expands our understanding of the molecular links between pigmentation levels and angiogenesis, and offers further avenues for closing the racial disparity gap in angiogenesis-dependent diseases.

## Acknowledgements

We thank Prof. Jerry Eichler for fruitful discussions. We are grateful to Prof. Dorothy C Bennett and Prof. Hyejung Jung for their generous gifts of MC1R cellular systems. This work was supported by US National Institutes of Health grant NEI-5-RO1-EY024046 to IA. AE was supported by the Israeli Ministry of Science and Technology (grant no. 17708).

## Declarations

### Ethics approval

The Boston Children’s Hospital (BCH) Institutional Review Board (IRB) has determined that this research qualifies as exempt from the requirements of human subject protection regulations.

### Availability of data and material

Gene expression data from the GTEx v7 release are available in dbGaP (dbGaP accession phs000424.v7.p2; https://www.ncbi.nlm.nih.gov/projects/gap/cgi-bin/study.cgi?study_id=phs000424.v7.p2).

### Code availability

All scripts used to analyze the data are available from the authors upon request.

### Authors’ contributions

IA and AE designed the study. SF carried out all gene expression analyses. MH performed all experiments. AE and IA interpreted the results and wrote the manuscript.

